# Stress granule transcript contrasts between colorectal cancer compartments depend on scoring and depth

**DOI:** 10.64898/2026.11.15.766839

**Authors:** Zaili Yang, Fei Li, Yin Chen, Yali Shi, Yu Li, Jinglin Wang

**Affiliations:** Department of Digestive Disease, Guizhou Provincial People’s Hospital, Guiyang, Guizhou, China

**Keywords:** stress granules, colorectal cancer, single-cell RNA sequencing, gene-set scoring, UCell, pseudobulk, sequencing depth, donor-level analysis, cancer-associated fibroblasts

## Abstract

Stress granule (SG) gene-set scores summarize transcription associated with stress-response machinery, but their stability across scoring definitions is uncertain. We compared fibroblast–epithelial expression contrasts for four fixed SG lists in two colorectal cancer single-cell cohorts containing 195,527 cells from 85 donors. Analyses used donor pairing, gene-wise standardized expression, within-cell ranks and raw-count pseudobulk summaries, with tissue and sequencing-depth sensitivities. At least 20 tumor cells per compartment were available for 15 and 29 paired donors. Pseudobulk SG-A contrasts were negative in both cohorts: median differences in mean log2 counts per million were −0.383 (95% confidence interval −0.493 to −0.319) and −0.493 (−0.614 to −0.384). Rank-based SG-B contrasts were also negative in both cohorts. In a post hoc comparison using identical depth-retained cells and weights, the second cohort had positive gene-Z estimates and negative rank-based estimates; the positive estimates did not meet the eight-test false-discovery threshold. Spatial comparisons depended on the fibroblast reference subtype, and the CosMx panel measured only 11 of 27 SG-B genes. RNA–protein correlations varied across genes, while organoid depletion calls were more frequent for selected translation-associated members than for the four core members. These findings establish reproducible relative-expression contrasts under specified definitions and show that their direction can change across scoring and depth conditions. Reporting the scoring definition, retained donor population, depth sensitivity and gene coverage together identifies the contrasts that reproduce and provides a defined basis for testing their relationship to SG formation.

## Introduction

Stress granules are dynamic assemblies of RNA and proteins that form under particular stress conditions. Their composition and assembly depend on cellular context, and the abundance of transcripts encoding SG-associated proteins is distinct from the physical presence or function of granules [1,2,3]. SG-associated processes have been implicated in cancer stress responses [4]. This biological interest has encouraged the use of transcriptional signatures, including an SG-related prognostic model previously reported for colorectal cancer (CRC) [5]. Such models address associations with clinical outcomes; they do not establish whether a gene-set score measures granule formation in individual cells.

Single-cell RNA sequencing can locate transcriptional programs within the tumor microenvironment, but comparisons between cell types depend on library size, gene detection and the expression characteristics of the selected genes. Method comparisons have documented limitations when bulk-derived signature scores are applied to single-cell data [6,7], and pseudobulk benchmarking has highlighted dependence on scoring design [8]. These findings motivate a specific question: how stable are fixed SG fibroblast–epithelial contrasts in CRC across scoring definitions, tissue selection and sequencing-depth sensitivities?

CRC atlases provide independent cohorts in which to examine this question [9,10]. We focused on epithelial cells and fibroblasts because their transcript abundance distributions differ and because fibroblast-associated programs may be relevant to tumor stress biology. We compared four nested SG gene sets with senescence-associated, ribosomal, housekeeping and proliferation-related reference measures. The reference measures were included to place the SG results in context, not to certify granule activity or senescence.

We reanalyzed two CRC single-cell datasets using donor-paired gene-Z, UCell and pseudobulk summaries. Sorted fractions and spatial data provided additional compartment comparisons. RNA–protein correspondence and organoid growth dependency supplied molecular and functional context for interpreting the selected genes. The study addresses the robustness of relative-expression contrasts and the biological interpretation supported by each measurement.

## Materials and methods

### Study design, data provenance and analysis units

This exploratory retrospective study reanalyzed processed public single-cell RNA sequencing (scRNA-seq) data from GSE132465 and GSE178341 [9,10] and examined independent measurement contexts described below. No new participants or specimens were collected. GSE132465 was analyzed from a processed Seurat object, and GSE178341 from an H5 count matrix with accompanying metadata. Source files were read without modification. S1–S6 Data provide gene definitions, cohort and donor records, numerical results and supporting inputs; S2 records processed-input hashes. Analysis scripts and source-verification records accompany the public reproducibility archive. The study was not preregistered. Tumor-focused analyses and depth sensitivities developed during exploratory analysis. UCell and pseudobulk specifications were recorded before those analyses were run, after the initial gene-Z results were known; the matched-cell diagnostic was added after score discordance was observed. Sample sizes followed data availability and the eligibility thresholds; no a priori power calculation was performed. GSE132465 contributed 18,539 epithelial cells and 3,462 fibroblasts from 23 donors. Tumor tissue contributed 17,469 epithelial cells and 1,501 fibroblasts, with both compartments represented in all 23 donors. GSE178341 contributed 168,295 epithelial cells and 5,231 fibroblasts from 62 donors. Tumor tissue contributed 113,216 epithelial cells from 62 donors and 3,031 fibroblasts from 58 donors, yielding 58 paired donors. Normal-tissue comparisons included 10 and 35 paired donors, respectively (Table 1; S2 Data). S2 Data records donor-by-tissue cell counts, the original patient–tissue mapping and eligibility under the one-cell and 20-cell thresholds. The two cohorts were analyzed separately; no pooled estimate across cohorts was calculated.

**Table 1.** Analysis populations and depth. GSE178341 tumor epithelial cells represent 62 donors; four lack paired fibroblasts. Normal and tumor donor counts overlap and must not be added. UMI summaries describe all selected cells, not only paired donors. Unique molecular identifier (UMI); epi, epithelial; fibro, fibroblast.

| **Cohort** | **Tissue** | **Epithelial cells** | **Fibroblasts** | **Paired donors** | **Epi UMI median** | **Fibro UMI median** |
|---|---|---|---|---|---|---|
| GSE132465 | Tumor | 17469 | 1501 | 23 | 19351 | 11301 |
| GSE132465 | Normal | 1070 | 1961 | 10 | 4519 | 6104 |
| GSE178341 | Tumor | 113216 | 3031 | 58 | 15261 | 9081 |
| GSE178341 | Normal | 55079 | 2200 | 35 | 7112 | 3680 |

Original processed cell annotations were retained. In GSE178341, PID identified donors, whereas PatientTypeID encoded patient–tissue samples and was not treated as an independent patient identifier. All cells from the same donor within a selected tissue and compartment were aggregated for the donor-level comparison. Cells therefore did not constitute independent inferential replicates. This distinction is necessary to avoid pseudoreplication in multi-donor single-cell studies [11,12]. The baseline comparison required at least one cell in each compartment; a minimum of 20 cells per donor and compartment was examined separately.

### Expression processing and gene definitions

For GSE132465, the supplied object contained an RNA assay and a recorded RNA normalization command. It contained no dimensional reductions or integrated expression assay. The target expression matrix was verified against the object’s RNA data layer, with maximum absolute difference below 10⁻¹⁵. Count-column sums agreed with the stored library sizes, and sampled non-zero entries agreed with log(1 + 10,000 × count/library size). Thus, the present scores used library-size-normalized, log-transformed RNA expression; no new Harmony integration was performed.

For GSE178341, the local H5 matrix contained 43,113 features and 370,115 cells. Every extracted target-gene count and every stored cell library size was checked against this H5 source, with zero discrepancies. This local matrix size differs from the 371,223 cells described in the GEO study summary; our analyzed subset is defined by the supplied matrix and metadata, rather than by that study-wide count. Target counts were transformed as log(1 + 10,000 × count/total UMI). The H5 feature metadata specified GRCh37_liftover_v28. No raw-read alignment, new cell filtering, doublet removal, ambient-RNA correction or cell-type reannotation was performed. Residual effects of these upstream steps remain limitations.

Four fixed, nested, author-curated SG-associated lists were retained from the existing analysis: SG-A (12 genes), SG-B (27 genes), core (4 genes) and nucleators (8 genes). SG-A was the principal list of interest and SG-B its extension; core contains G3BP1, G3BP2, TIA1 and TIAL1. The historical list names identify these working definitions rather than externally trained activity signatures or a uniform functional classification. Exact membership, overlap and cohort coverage are provided in S1 Data. SG literature provides context for the included RNA-binding, translation-associated and assembly-related proteins [1,3]. The lists were not derived from the catalog comparisons below, and no granule-imaging calibration was used.

For a retrospective annotation check, all 27 SG-B members were mapped to three resources: direct, non-NOT human GOA annotations to GO:0010494 (cytoplasmic stress granule); the source classifications in the processed U2OS cuffdiff table from Khong et al. (GSE99304) [13]; and the SG protein-profile table of Hu et al. [14]. The retained GOA file was generated on 28 July 2026 using the 26 July 2026 GO release and was accessed for this check on 5 October 2026. S3 Table records exact list membership, resource version, source labels and gene-level evidence: 22/27 genes had the specified GO annotation, and 16/27 were represented in the Hu profile table, all labeled there as previously known SG proteins. In the Khong table, ATXN2, EIF4G1 and PRRC2C were classified as enriched, EIF4E as depleted, and the remaining 23 as neither. Protein association with SGs and enrichment of the corresponding mRNA are distinct annotations. These overlaps describe the fixed lists against public resources; they neither reconstruct the original selection process nor validate a transcript score as granule activity.

Reference measures comprised the published 125-gene SenMayo set [15], a 25-gene senescence-associated secretory phenotype (SASP) subset, 15 ribosomal genes, six genes labeled housekeeping in the source analysis, and MKI67 expression. Non-gene header tokens were removed from the supplied SenMayo definition. Of its canonical genes, 123 were available in GSE132465 and 124 in GSE178341; BEX3 and CCL3L1 were absent from the former target matrix, and PECAM1 was absent from the latter source feature list. A sensitivity analysis used the 122 genes common to both datasets. All other defined signature genes were available. SASP is fully nested within the SenMayo list and was not considered independent validation. The housekeeping list includes VIM and B2M, so it was treated as an empirical reference rather than a biologically invariant technical control.

### Scores and donor-level contrasts

Within each cohort, each gene’s log-normalized expression was centered and scaled using all analyzed epithelial cells and fibroblasts, including tumor and normal tissue. The score was the arithmetic mean of these gene-wise Z scores over available, non-constant genes. The scaling reference was held fixed across tissue and depth sensitivity analyses. This cell-weighted reference facilitates comparison within a cohort but is influenced by compartment and tissue composition; it is not an externally calibrated measurement scale. MKI67 was reported as unstandardized log-normalized expression in both cohorts.

For each donor, we calculated the mean score in fibroblasts minus the mean score in epithelial cells. The reported effect was the median of these donor differences, giving donors equal weight in the final summary. A positive contrast indicates higher expression of the selected gene set in the retained fibroblasts; it does not establish greater SG assembly. Nonparametric percentile confidence intervals were obtained by resampling donors with replacement 10,000 times. Two-sided Wilcoxon signed-rank tests were performed using SciPy’s automatic method selection.

These tests concern the signed-rank location null and require appropriate symmetry for a simple location interpretation. Their P values are not inversions of the percentile confidence intervals for the median; disagreement between a confidence interval and a test threshold can therefore occur. Benjamini–Hochberg adjustment was applied across the nine measures within each cohort and each analysis specification. These q values do not control a single error rate over all explored tissues, constructions, thresholds and sensitivity analyses. Effects and their uncertainty were emphasized over threshold-based classification. No equivalence margin was specified, and no non-significant contrast was interpreted as evidence that a biologically relevant effect had been excluded [16]. Standardized score magnitudes were not interpreted as directly comparable biological effect sizes across different gene sets.

Multiplicity was controlled within each analysis specification. Gene-Z comparisons used nine measures per cohort; UCell and set-level pseudobulk comparisons used four sets across two cohorts (eight tests); and FAP comparisons used nine measures per cohort per specification, yielding 36 reported rows. Sorted-fraction tests used four SG sets per construction. GeoMx region and paired-compartment analyses used separate eight-test families. CosMx and RNA–protein analyses used 11 and 27 genes, respectively, per specification. edgeR adjustment covered all filtered genes within each cohort. Organoid intervals were descriptive and pointwise. These local adjustments do not control a project-wide error rate.

### Depth dependence, detection and balancing

Depth was represented by log₁₀(total UMI). Spearman correlations between depth and scores were calculated descriptively in pooled cells and separately in epithelial cells and fibroblasts, both across all tissues and within tissue subsets. Because cells were clustered within donors, these correlations were not used for cell-level significance tests. Mean per-gene detection was the number of non-zero gene–cell entries divided by the product of the number of available genes and cells. The all-zero fraction was the proportion of cells with no detected member of a given set. These two quantities answer different questions and were reported separately.

The principal depth sensitivity analysis divided log₁₀ UMI into fixed-width 0.15 bins within each donor. A bin was retained only if it contained at least five epithelial cells and five fibroblasts. Within-bin mean-score differences were weighted by the smaller compartment cell count and averaged to obtain one difference per donor. Donors without an eligible bin were excluded. This procedure reduces observed depth imbalance but changes both the cells and, potentially, donors represented. It is therefore a depth-balanced sensitivity analysis rather than a causal correction for technical noise.

To distinguish donor selection from within-donor changes, we repeated the unbalanced comparison using precisely the donors retained by depth balancing. We also inspected residual log-UMI imbalance and varied bin width to 0.10 and 0.20. Additional analyses restricted each cohort–tissue subset to the intersection of compartment-specific 5th–95th percentile depth intervals. This common-window restriction was not labeled matching because it does not enforce equal depth distributions. All-tissue, normal-only and sample-paired summaries were retained as secondary analyses. For the latter, sample-specific contrasts were averaged within donors before summarizing donors.

### Alternative constructions and FAP-associated comparisons

For each gene set, we compared the primary mean gene-Z score with mean log-normalized expression, the number of detected genes, mean expression of the five most highly expressed set members in each cell, and the mean of gene-wise depth-residual Z scores. Residual scores were obtained by fitting a linear expression–log-UMI relationship for each gene across the two analyzed compartments and tissues within a cohort. The top-five summary is a custom expression statistic, not an implementation or approximation of UCell or AUCell. These constructions have different scales and estimands; their signs were compared without interpreting ratios of their magnitudes as technical inflation. UCell is a distinct rank-based scoring procedure [17]; its separately specified execution is described below.

Within tumor fibroblasts, an exploratory fibroblast activation protein (FAP) analysis compared cells with detected FAP expression against cells with zero measured FAP expression. FAP has established relevance to tumor-associated stroma [18], but failure to detect its transcript does not establish protein negativity. The unbalanced analysis required at least five cells in each FAP group per donor. Depth balancing used the same 0.15 bins but required at least three cells in each group per bin. Inference again used donors, not pooled cells, as the sampling units. Each donor contributed one FAP-detected minus FAP-undetected contrast; donor resampling supplied the 95% percentile interval, and BH adjustment covered nine measures within each cohort and FAP analysis specification. S5 Data supplies the fibroblast input scores, FAP detection labels, donor eligibility, depth-bin counts and weights, donor differences and all 36 summary rows. This aggregation-based treatment of patient clustering did not fit a random-intercept model.

### Competitive random-gene-set analyses

Competitive analyses were performed only in GSE132465, where the supplied all-gene matrix and candidate pools were available. The universe contained 25,258 non-constant genes. For each target set, 2,000 random sets of equal size were sampled either from the whole universe or from the stored 500-candidate pools matched on standardized mean expression, log standard deviation and detection frequency. The matched pools excluded the selected target and control genes; the unmatched universe retained all eligible genes. Genes were sampled without replacement within a draw. Pools were defined using the full epithelial–fibroblast analysis matrix and were reused for the tumor-only analysis; they were not re-optimized for tumor tissue. The historical processed project contains the pool definitions; its pool-map hash is provided in S2. S4 Data supplies all 2,000 resulting draws per set, tissue and null specification and the equal-tail summaries. The candidate pool map and all-gene matrix are not embedded in S1–S6, so new random draws require those original processed inputs.

Each random set was scored with the same gene-wise scaling and donor aggregation as the target. Competitive two-sided tail areas were calculated as twice the smaller empirical tail, with a plus- one correction, capped at one. We did not center the null distribution at zero: random gene sets can themselves differ between compartments. These analyses compare a target set with alternative sets conditional on the observed donors [19,20]. They do not estimate donor-sampling power, supply an equivalence bound or establish the minimum biological effect that could be excluded. Both matched and unmatched distributions were reported, without selecting one according to which produced a preferred result.

### Full-gene UCell analysis

UCell and pseudobulk specifications were recorded after the initial gene-Z results were available and before these additional analyses were run. Both analyses required at least 20 tumor cells per donor in each compartment. The four fixed SG lists were retained, with SG-A as the principal construct of interest. No additional genes or methods were selected according to significance. These analyses are exploratory robustness comparisons, not independent confirmation of a preregistered hypothesis.

UCell scores [17] were calculated from the full raw-count gene background using unmodified official core functions, data_to_ranks_data_table and u_stat_signature_list, from CarmonaLab/UCell commit f778e091f19e28d3d05bde134663fd6b2f938267 (DESCRIPTION version 2.17.1). The core source was called through a sequential chunk adapter pinned to the specified commit, without changing the scoring functions. Parameters were maxRank = 1500, average ties and zero imputation for absent signature genes, with no smoothing. Duplicate feature symbols were summed before ranking. GSE132465 had 25,655 unique gene symbols; GSE178341 had 43,113 features mapping to 41,195 unique symbols. All 27 SG genes had one source feature each in GSE178341, and all four signatures were fully represented in both cohorts. Within-cell rank calculations used all genes, not only signature members. Official-core results were checked against an independent rank formula with ties, absent genes and cell-subset invariance.

Scores were averaged within donor and compartment, followed by the same median donor contrast, bootstrap interval and signed-rank test described above. The main UCell family comprised four signatures across two cohorts (eight BH-adjusted tests). Sensitivities used the one-cell minimum, or depth balancing among min20 donors, with an additional comparison of the same donors before balancing. Each sensitivity specification constituted a separate eight-test family. Correlations with UMI were descriptive. After observing a difference in direction between UCell and the original gene-Z depth analyses, we added a diagnostic comparison using exactly the UCell-retained donors, cells, bins and weights for both scores. This comparison was not prespecified; it was reported as an eight-test family and logged as a post-result diagnostic.

### Paired raw-count pseudobulk analysis

Raw integer counts were summed by donor and tumor compartment. Library sums were reconciled with the constituent cells, and the main design retained donors meeting the 20-cell minimum in both compartments. edgeR 4.10.1 [21] was used with filterByExpr, trimmed mean of M values (TMM) normalization, robust dispersion estimation and robust quasi-likelihood fitting. The design was ∼ PID + compartment, with epithelial cells as the reference. Model rank and residual degrees of freedom were checked. Gene-level BH adjustment covered all retained genes within each cohort, and all 27 SG-B genes were reported regardless of significance.

To describe a set-level quantity on the same donor samples, we averaged TMM log2-CPM (prior.count = 2) over each fixed gene list, using effective library sizes from the fitted count pipeline. These summaries retained all available set members and were not supplied as inputs to the count model. Their paired median contrasts, donor-bootstrap intervals and signed-rank P values were reported as a separate eight-test BH family. A pseudobulk average of log-CPM, a cell-averaged UCell rank score and a cell-weighted gene-Z score are different estimands. Aggregation also weights cells by their contribution to the count pool; TMM measures relative RNA abundance and does not recover absolute transcripts per cell. Neither approach identifies granule activity.

### Computation and reproducibility

Reanalysis used Python 3.12.14, NumPy 2.3.5, pandas 3.0.1 and SciPy 1.16.2, with h5py for source verification. The main random seed was 20260926; separate deterministic seeds for matched-donor and additional sensitivity analyses are recorded in the scripts. The Seurat object was inspected in R 4.6.1 with Matrix and SeuratObject. The extension used official UCell core source, edgeR 4.10.1, limma 3.68.4 and locfit 1.5-9.12; execution records include source hashes and the frozen analysis plan. Output tables were used directly to generate manuscript tables and figures. S3 Data reports gene-Z effects, confidence intervals, donor differences and tissue/depth sensitivities; S4 reports detection, alternative constructions and competitive backgrounds; S6 reports all UCell and pseudobulk specifications, gene-level edgeR results, filtering and model-sample metadata. S5 and S6 include scripts for recomputing their summary statistics from the supplied inputs. Each of S1–S6 includes a file inventory and a column-level data dictionary. Source-level checks validated the supplied processed inputs; they did not reproduce the original studies from raw reads.

### External sorted and spatial transcriptomic analyses

GSE39396 supplied 24 processed microarrays from six patients with paired FAP-positive, EPCAM-positive, CD31-positive and CD45-positive fractions [22]. We mapped GPL13158 probes to genes and averaged log2 probe values for the main analysis, with median-probe and gene-Z sensitivities. The inferential contrast was each patient’s mean FAP-minus-EPCAM set expression; exact sign permutations and 10,000 patient bootstraps described uncertainty, with BH correction across four SG sets. Protein-based sorting supported fraction identity but did not measure granules.

The published early-CRC resource [23] supplied GeoMx and CosMx data through Zenodo record 17671259. The GeoMx whole-transcriptome atlas (WTA) experiment contained 281 retained segments from ten patients. We excluded the T3 specimen from T1 comparisons, averaged eligible region-of-interest (ROI) contrasts within patients, and compared invasive front with core separately in PanCK-positive and PanCK-negative segments. Nine T1 patients contributed region comparisons and eight contributed paired compartment comparisons. Mean log2(q_norm + 1) expression was primary; author batch-corrected log expression was a sensitivity. The cancer transcriptome atlas (CTA) experiment had no overlap with the 27 SG genes and was excluded from SG scoring. PanCK-negative segments were mixed stroma. Patient sign permutations and bootstrap intervals were used; the eight region tests combined four sets and two compartments. Reference-list analyses were added after inspecting SG results and were descriptive.

For CosMx experiment 2, the complete archive was checked against the author MD5 before expression extraction. We retained the source study’s quality control (QC) and cell annotations, joined by unique cell identifiers, and audited the coverage of the fixed lists. All four lists failed the prospectively specified 80% panel-coverage gate; complete SG scores were therefore not calculated. Before inspecting target gene effects, we amended the analysis to test every measured SG-B member individually using log(1 + 10,000 × raw count/total measured-panel count). These gene measurements were not renamed as an SG activity signature. Coverage, the amended contract and every result are supplied in S7 Data. Trophocyte-like cancer-associated fibroblasts (CAFs) were compared with trophocytes within patients having at least 20 cells of each label. Telocytes were a secondary comparator. A label-confidence sensitivity required prediction.score.max >0.6 and the same 20-cell minimum; the author k8 clusters were described without relabeling them according to the target genes. Depth sensitivity used five pooled depth quantiles within each patient/comparison, retaining bins with at least five cells of each label and weighting by the smaller group count; at least 20 cells of each label had to remain. We separately report raw contrasts for these same retained patients.

We reused the published 50-µm neighbor-composition table, rather than reconstructing radius neighborhoods with different QC rules. CAF exposure was the sum of trophocyte-like CAF and MMP CAF fractions; the table did not contain a telocyte-like CAF column. Zero-sum neighborhood rows were excluded. For each eligible patient, standardized epithelial gene expression was regressed on standardized CAF fraction and log panel counts with field-of-view (FOV) fixed effects. At least 100 epithelial cells and nonzero residual exposure variation were required. FOV identifiers combined slide and FOV; models were fitted separately per patient. Sensitivities added epithelial-state indicators, restricted to CRC/oncofetal epithelium, used non-SG panel counts for depth, or replaced CAF exposure with the seven annotated fibroblast-related neighbor categories. Absolute neighbor counts were unavailable and were not adjusted for. We summarized patient slopes with equal weight, 10,000 patient bootstraps, exhaustive two-sided sign permutations and 11-gene BH adjustment within each specification. No cell-level P values were used. The sign-permutation interpretation assumes symmetry/exchangeability of patient effects under the null; percentile intervals and discrete permutation tests are not exact inversions. Cell labels came from author transcriptomic clustering/label transfer, so these are exploratory label-conditioned associations with possible feature overlap, not independent validation of CAF identity or granule activity. Cross-sectional stage composition was descriptive. The analysis specification and metadata-based refinements were saved before viewing target expression results, but were not externally preregistered.

### RNA protein correspondence and external functional context

We matched 96 tumor identifiers between the prospective CPTAC-COAD tumor proteome and RNA matrices distributed through LinkedOmics [24,25]. The source tables contained 8067 protein rows across 97 tumors and 13482 RNA rows across 106 tumors; RNA was RSEM upper-quartile log2(value + 1), whereas protein values were tandem mass tag (TMT) log ratios. Spearman correlations for the fixed 27 genes used complete pairs per gene, 10,000 patient bootstraps and 27-gene BH adjustment. Rank residualization against expression-derived tumor purity was a sensitivity. This bulk analysis concerned total protein abundance, not CAF-specific proteins or granule recruitment. We annotated SG members against published U2OS heat-stress/recovery protein profiles [14]; only published profile means were used, so we did not create replicate-level dynamic tests or CRC-specific temporal claims.

The published organoid CRISPR resource [26] was analyzed using the author’s quality ordering to retain one screen per model. Of 85 colorectal models, two adenomas were excluded, leaving 83 malignant models from 81 patients; 24 of 27 SG genes were assayed. Author BAGEL2 depletion calls were retained, and CRISPRcleanR scaled log-fold changes were summarized after averaging models from the same patient and then taking the median across patients. Patient-bootstrap intervals used 10,000 draws. These descriptive intervals were not simultaneous intervals across genes, and no new hypothesis-test family was declared. Missing genes were not assigned zero dependency, and a Chronos threshold was not applied. Duplicate-library concordance was assessed descriptively. The related Human Cancer Models Initiative (HCMI) resource [27] was audited for cohort overlap and clinical feasibility; insufficient outcome events or missing survival times precluded fitting a prognostic model. No independent validation claim was made between overlapping organoid resources.

### Integration across measurement contexts

The core question remained the interpretation of fixed SG transcript lists across measurement contexts. We retained spatial/subtype expression as direct context for the compartment analysis, and used sorted fractions, bulk RNA–protein correspondence and organoid dependency as secondary evidence with distinct endpoints. Published non-CRC protein dynamics supplied biological context only. No evidence score or meta-analysis combined unlike endpoints. All follow-on analyses were exploratory, with multiplicity controlled locally rather than across the entire project. Complete null, positive and heterogeneous results are supplied in S7–S8 Data.

### Ethics approval and consent to participate

This study reanalyzed publicly available, processed, de-identified data from previously published studies and involved no new participant recruitment, contact or specimen collection. Guizhou Provincial People’s Hospital does not conduct institutional ethics review or issue ethics exemption determinations for bioinformatics-only studies of this type. No institutional ethics approval or exemption was issued for this secondary analysis. The original studies describe their ethics approvals and consent procedures [9,10,22,23,24,26,27]. Consent for publication of identifiable participant information is not applicable. None of the six authors had access to information that could identify individual participants during or after this study.

Research-access dates were reconstructed from retained execution records. Local access to processed GSE132465 and GSE178341 tables is documented on 20 September 2026. Access for sorted-fraction, spatial, RNA–protein and organoid analyses is documented on 26 September 2026. The retained gene-catalog files were accessed and verified for this revision on 5 October 2026. These are documented analysis and verification access dates, not the original download dates or the dates of participant recruitment in the source studies.

Clinical trial number: not applicable.

### Use of AI-assisted tools

OpenAI Codex assisted with language editing, figure polishing and supporting-file preparation. Numerical results came from scripts executed against the specified data. Checks included processed-input reconciliation, patient-summary reconstruction, an independent UCell rank formula, comparison of manuscript estimates with saved results, and direct inspection of the gene-catalog source rows. Figures were plotted from recorded numerical outputs. The authors reviewed the outputs and remain responsible for the scientific content, interpretations and declarations.

## Results

### Gene-Z contrasts varied across cohorts and tissues

The two datasets contained 195,527 epithelial cells and fibroblasts from 85 donors, of whom 81 had paired tumor compartments (Table 1). We report gene-Z contrasts and depth sensitivities, followed by rank-based UCell and pseudobulk comparisons with stricter cell eligibility. The median tumor-cell library size was higher in epithelial cells than fibroblasts in both cohorts: 19,351 versus 11,301 UMI in GSE132465 and 15,261 versus 9,081 UMI in GSE178341. The normal-tissue pattern differed between cohorts, supporting separate tissue summaries.

For tumor-only SG-A, the median fibroblast–epithelial contrast was −0.066 (95% confidence interval −0.140 to 0.039; q = 0.214) in GSE132465 and −0.042 (−0.081 to 0.001; q = 0.064) in GSE178341. SG-B was also negative: −0.010 (−0.156 to 0.018; q = 0.264) and −0.057 (−0.091 to −0.012; q = 0.017), respectively. Core and nucleator contrasts were negative in the unbalanced tumor comparisons, but their uncertainty and signed-rank results did not support a common, well-resolved effect across cohorts (Table 2; Fig 1). Thus, the results were not uniformly null, and they did not show a consistent unbalanced fibroblast enrichment.

**Table 2.**
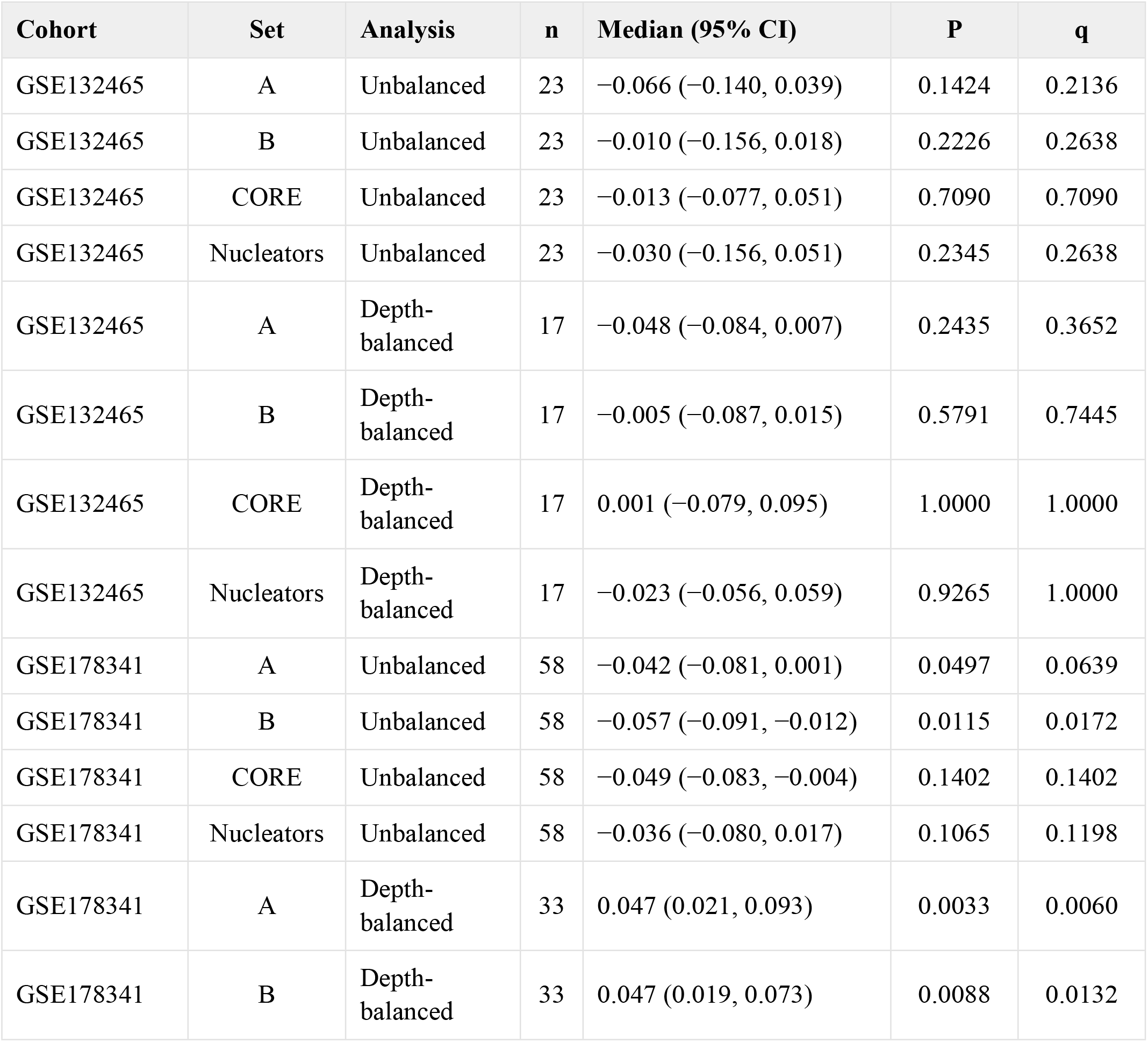
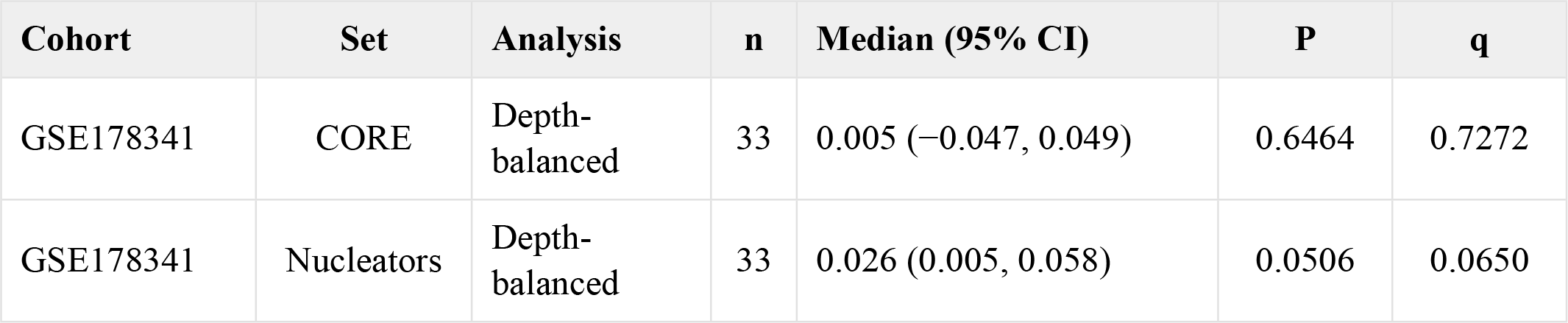
Tumor SG contrasts and uncertainty. . Difference = fibroblast minus epithelial mean score within donor, summarized by its median across donors. Confidence intervals use donor bootstrap percentiles; P values use two-sided Wilcoxon signed-rank tests. q values adjust nine measures within each cohort and specification. The interval and test target different inferential quantities.

| Cohort | Set | Analysis | n | Median (95% CI) | P | q |
|---|---|---|---|---|---|---|
| GSE132465 | A | Unbalanced | 23 | −0.066 (−0.140, 0.039) | 0.1424 | 0.2136 |
| GSE132465 | B | Unbalanced | 23 | −0.010 (−0.156, 0.018) | 0.2226 | 0.2638 |
| GSE132465 | CORE | Unbalanced | 23 | −0.013 (−0.077, 0.051) | 0.7090 | 0.7090 |
| GSE132465 | Nucleators | Unbalanced | 23 | −0.030 (−0.156, 0.051) | 0.2345 | 0.2638 |
| GSE132465 | A | Depth-balanced | 17 | −0.048 (−0.084, 0.007) | 0.2435 | 0.3652 |
| GSE132465 | B | Depth-balanced | 17 | −0.005 (−0.087, 0.015) | 0.5791 | 0.7445 |
| GSE132465 | CORE | Depth-balanced | 17 | 0.001 (−0.079, 0.095) | 1.0000 | 1.0000 |
| GSE132465 | Nucleators | Depth-balanced | 17 | −0.023 (−0.056, 0.059) | 0.9265 | 1.0000 |
| GSE178341 | A | Unbalanced | 58 | −0.042 (−0.081, 0.001) | 0.0497 | 0.0639 |
| GSE178341 | B | Unbalanced | 58 | −0.057 (−0.091, −0.012) | 0.0115 | 0.0172 |
| GSE178341 | CORE | Unbalanced | 58 | −0.049 (−0.083, −0.004) | 0.1402 | 0.1402 |
| GSE178341 | Nucleators | Unbalanced | 58 | −0.036 (−0.080, 0.017) | 0.1065 | 0.1198 |
| GSE178341 | A | Depth-balanced | 33 | 0.047 (0.021, 0.093) | 0.0033 | 0.0060 |
| GSE178341 | B | Depth-balanced | 33 | 0.047 (0.019, 0.073) | 0.0088 | 0.0132 |
| Cohort | Set | Analysis | n | Median (95% CI) | P | q |
| GSE178341 | CORE | Depth-balanced | 33 | 0.005 (−0.047, 0.049) | 0.6464 | 0.7272 |
| GSE178341 | Nucleators | Depth-balanced | 33 | 0.026 (0.005, 0.058) | 0.0506 | 0.0650 |

**Fig 1.** Tumor-only SG gene-set contrasts before and after within-donor depth balancing. Panels a and b show GSE132465 and GSE178341, respectively. Points denote medians of donor-level fibroblast-minus-epithelial mean scores; bars are 95% percentile confidence intervals from 10,000 donor bootstraps. Blue points include all paired tumor donors (23 and 58); orange points include depth-balanced subsets (17 and 33). Gene numbers are shown in parentheses. The two colors therefore differ in both retained cells and donor inclusion; Fig 3 holds donor inclusion fixed. Effects are averages of gene-wise Z scores, not standardized biological effect sizes across sets.

Normal-only SG-A contrasts were 0.076 (95% CI 0.039 to 0.154; n = 10; q = 0.00352) in GSE132465 and −0.001 (−0.034 to 0.031; n = 35; q = 0.778) in GSE178341. All-tissue SG-A contrasts were −0.056 (−0.109 to 0.014; n = 23) and −0.038 (−0.074 to −0.010; n = 61), respectively. These secondary summaries used the same fixed gene-scaling reference as the tumor analysis (S2–S3 Data). The tissue-specific patterns are descriptive; no formal tissue-by-compartment interaction was tested.

SenMayo and SASP contrasts were positive in both cohorts. SenMayo median differences were 0.397 and 0.804, and SASP differences were 0.652 and 1.263. Using the 122 common SenMayo genes gave similar estimates of 0.398 and 0.807. Ribosomal and MKI67 differences were negative, whereas the housekeeping-list difference was positive (S1 Fig). These empirical reference lists showed compartment-associated expression differences under the same analysis.

### Depth associations persisted within cell compartments

Across all analyzed tissues, SG-A scores correlated with log-UMI in the pooled data (Spearman ρ = 0.400 and 0.606). Correlations remained substantial within epithelial cells (0.386 and 0.613) and fibroblasts (0.316 and 0.446) (Fig 2). Consequently, the pooled association could not be attributed solely to mixing the two compartments. Correlations within narrow depth bins are difficult to interpret because restricting the range of the predictor can attenuate them.

**Fig 2.**
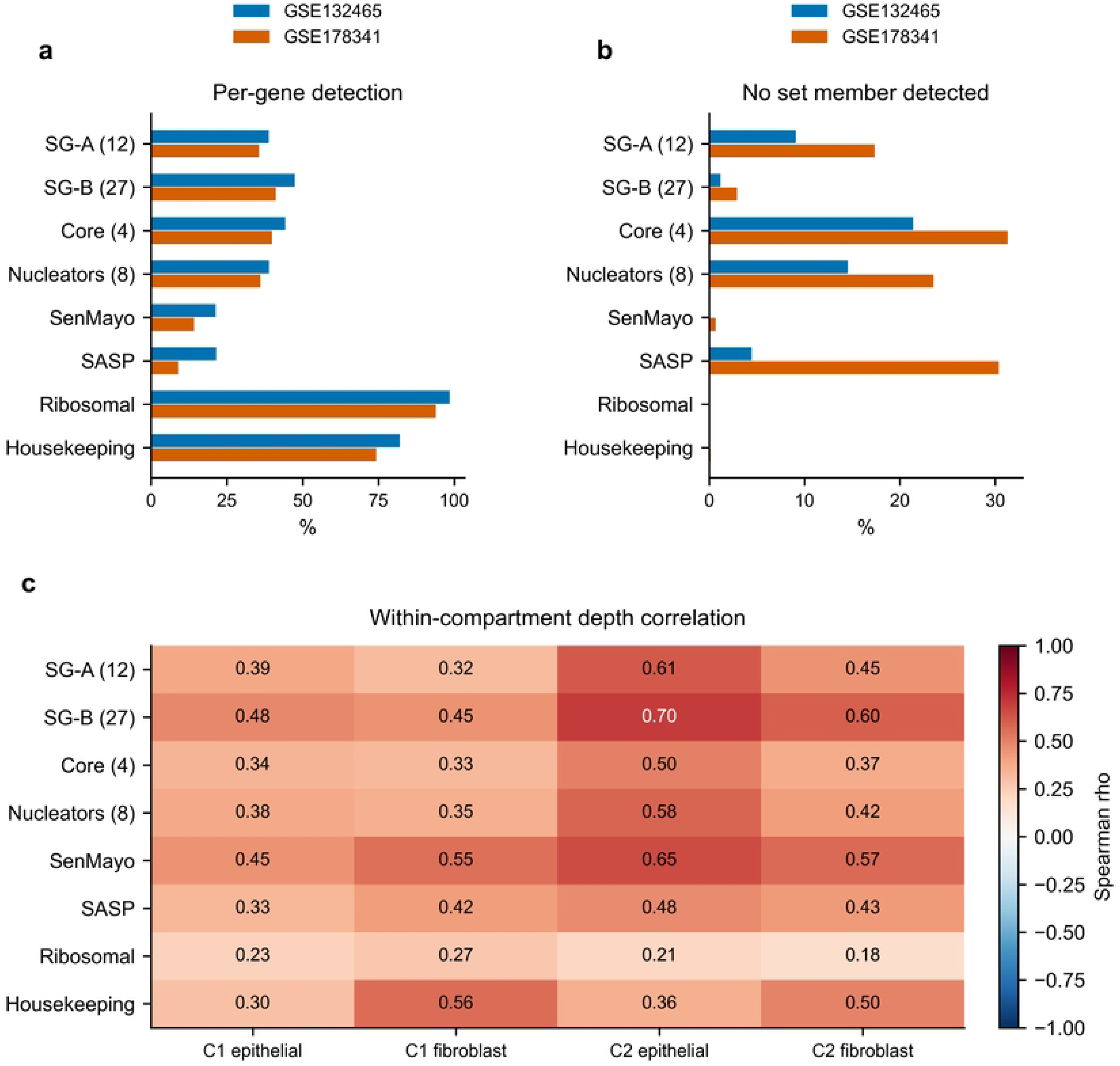
Gene detection and within-compartment depth associations. Panel a shows mean per-gene detection; panel b shows the fraction of cells with no detected member of the set. Both use all analyzed epithelial cells and fibroblasts across tumor and normal tissue (22,001 and 173,526 cells). Panel c shows descriptive Spearman correlations between score and log₁₀ UMI within each compartment. C1, GSE132465; C2, GSE178341; epi, epithelial cells; fibro, fibroblasts. Correlations are descriptive cell-level summaries, without cell-level significance tests. SG lists overlap, and the SASP list is contained within SenMayo.

Mean per-gene SG-A detection was 38.8% and 35.6% in the two cohorts; SG-B detection was 47.4% and 41.1%. SenMayo detection was lower, at 21.3% and 14.2%. In contrast, the proportion of cells with no detected member was 9.1% and 17.3% for SG-A, 1.2% and 2.9% for SG-B, and 21.4% and 31.3% for the four-gene core set (Fig 2; S4 Data). Thus, detection per gene and the probability of detecting any set member described different aspects of sparsity. Small sets had appreciable all-zero detection fractions despite higher per-gene detection than SenMayo.

### Depth balancing changed the second cohort’s contrasts

Within-donor depth balancing retained 17 of 23 paired tumor donors in GSE132465 and 33 of 58 in GSE178341. Among these donors, the median fibroblast–epithelial log-UMI difference was reduced from −0.155 to −0.004 in the first cohort and from −0.273 to −0.004 in the second.Residual imbalance was reduced rather than eliminated, and the excluded donors limit generalizability to the full cohort.

The depth-balanced SG-A contrast was −0.048 (−0.084 to 0.007; q = 0.365) in GSE132465 and 0.047 (0.021 to 0.093; q = 0.006) in GSE178341. Corresponding SG-B contrasts were −0.005 (−0.087 to 0.015; q = 0.745) and 0.047 (0.019 to 0.073; q = 0.013). SG-A and SG-B therefore retained positive depth-balanced estimates in the second cohort. Core estimates remained uncertain, and the positive nucleator estimate in that cohort had q = 0.065 (Table 2). Restricting the unbalanced analysis to the same retained donors preserved the change in direction in GSE178341: SG-A shifted from −0.037 to 0.047 and SG-B from −0.036 to 0.047 (Fig 3; S1 Table). The median within-donor changes were 0.075 and 0.091, respectively. In GSE132465, the corresponding SG-A contrast remained negative. Bin-width sensitivity analyses produced negative SG-A and SG-B estimates in the first cohort and positive estimates in the second across widths of 0.10, 0.15 and 0.20 (S3 Data). The cohort-specific direction was therefore stable across the three bin widths tested.

**Fig 3.**
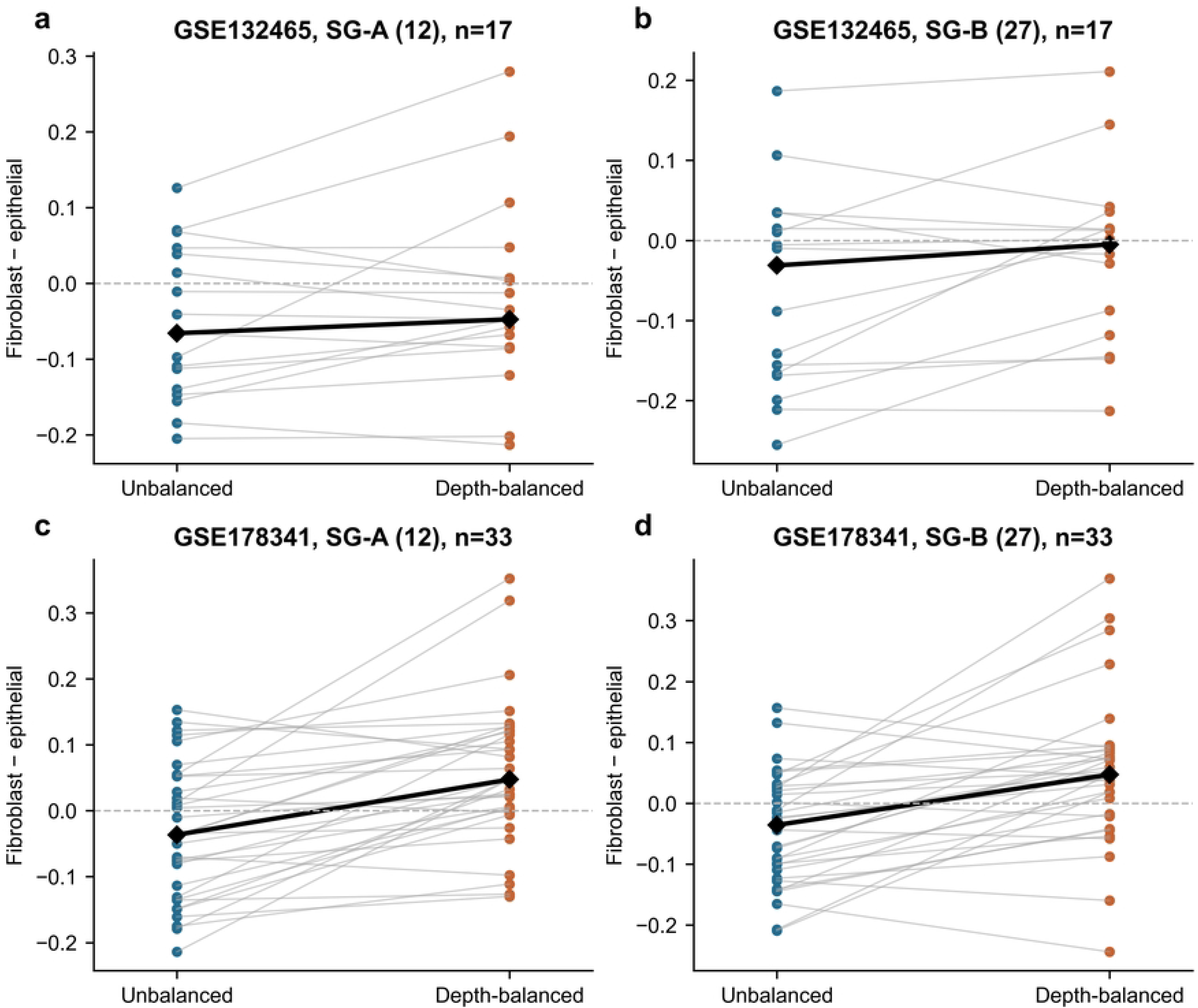
Paired changes among the same retained donors. SG-A and SG-B contrasts are shown for GSE132465 (a,b) and GSE178341 (c,d). Each thin line links one donor’s unbalanced and depth-balanced contrast; black diamonds denote the median across donors. Only donors retained by depth balancing are displayed (17 and 33). The selected cells still differ between the two analyses. Depth balancing used 0.15-wide log₁₀ UMI bins with at least five cells per compartment and weighted bins by the smaller compartment cell count.

### Score construction and depth affected SG and FAP contrasts

The sign of the tumor contrast depended on score construction for two of four SG lists in GSE132465 and all four in GSE178341 (S2 Fig). In the second cohort, depth-residual scores were positive for all four lists, whereas the primary gene-Z contrasts were negative. This comparison shows dependence on the analytical estimand. It does not identify a uniquely correct construction, because regression can remove both technical and biological components of RNA abundance.

Within tumor fibroblasts, unbalanced donor-level SG-A differences between FAP-detected and FAP-undetected cells were 0.080 (95% CI 0.045 to 0.171; n = 16; q = 0.00808) in GSE132465 and 0.179 (0.064 to 0.225; n = 29; q = 0.0000160) in GSE178341. Depth balancing retained 12 and 18 donors and gave estimates of −0.032 (−0.063 to 0.003; q = 0.277) and 0.021 (−0.081 to 0.082; q = 0.790), respectively (S5 Data). No SG measure met q < 0.05 in either depth-balanced FAP comparison. The reduced donor counts, sparse FAP measurement and differing retained cells prevent these results from establishing biological absence of an association. The donor-level input and eligibility tables distinguish patient clustering from the selection changes induced by depth balancing.

### SG contrasts were unexceptional against first-cohort competitive backgrounds

In tumor tissue from GSE132465, none of the four SG contrasts lay in an extreme tail of either competitive distribution (S3 Fig). For SG-A, the observed difference was −0.066, compared with a matched-null mean of −0.070 and an unmatched-null mean of 0.038; the respective equal-tail P values were 0.936 and 0.241. SG-B, core and nucleator matched P values were 0.349, 0.714 and 0.736. By contrast, all four multi-gene reference lists had matched competitive P ≤ 0.014. The matched null therefore did not uniformly fail to distinguish reference contrasts in this tumor-only analysis.

The SG contrasts in GSE132465 were typical of the specified alternative gene sets. This competitive comparison is conditional on the observed donors and addresses gene-set specificity rather than donor-sampling power. Full tissue, cell-count and common-window sensitivities are provided in S3 Data, with competitive results and draws in S4 Data.

### UCell and pseudobulk analyses measured distinct compartment contrasts

The extension required at least 20 tumor cells in each compartment, retaining 15 paired donors in GSE132465 and 29 in GSE178341. SG-A UCell contrasts were −0.007 (−0.015 to 0.001; q = 0.122) and −0.007 (−0.013 to −0.003; q = 0.00195), respectively. SG-B UCell contrasts were −0.014 (−0.024 to −0.007; q = 0.0164) and −0.018 (−0.031 to −0.014; q = 8.94e-08). The original gene-Z scores were also recomputed on these same donor sets so that the main method comparison was not driven by different donor eligibility (Fig 4).

**Fig 4.**
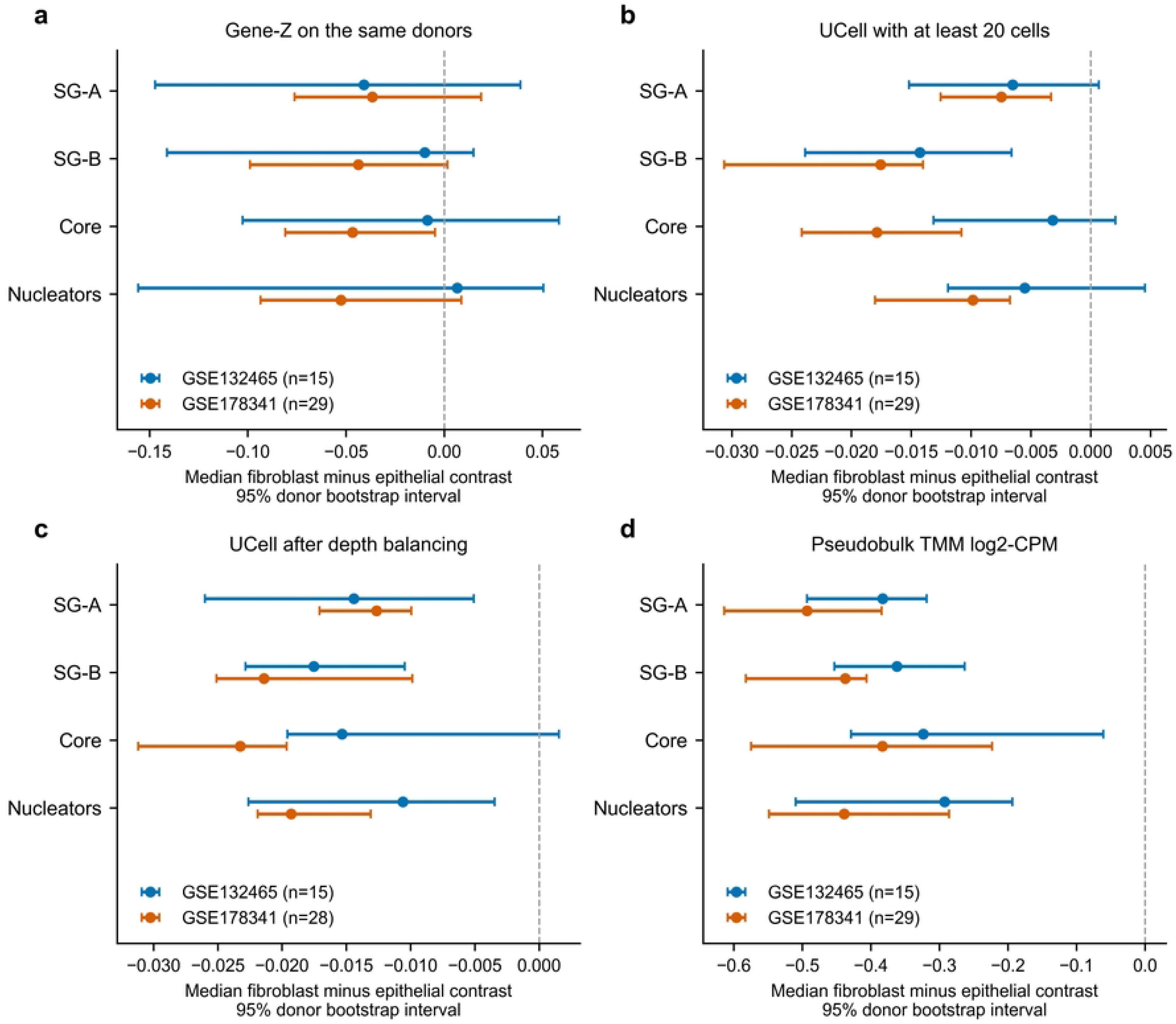
Compartment contrasts across scoring definitions. a, Gene-Z scores restricted to the same donors meeting the 20-cell minimum as the UCell and pseudobulk analyses. b, UCell scores calculated with official core functions (maxRank 1500, average ties, full gene background). c, UCell scores after within-donor depth balancing among eligible min20 donors; donor attrition is shown in each legend. d, Mean TMM log2-CPM over each fixed gene set after raw-count pseudobulk aggregation. Points show median donor differences, fibroblast minus epithelial; bars show 95% percentile confidence intervals from 10,000 donor bootstraps. Axes differ because the scores have different units. Full P and BH-adjusted q values are in S6 Data (supplement_results.csv); overlapping lists are not independent validations.

Depth-balanced UCell analysis retained 15 and 28 of these donors. SG-A contrasts were −0.014 (−0.026 to −0.005; q = 0.0201) and −0.013 (−0.017 to −0.010; q = 2.72e-06). S6 Data includes the unbalanced values for exactly these retained donors, as well as within-donor changes; donor selection and within-donor changes are not interchangeable explanations. The corresponding SG-A pseudobulk log-CPM contrasts were −0.383 (−0.493 to −0.319; q = 9.77e-05) and −0.493 (−0.614 to −0.384; q = 9.93e-09). For SG-B, the estimates were −0.362 (−0.453 to −0.263; q = 0.00014) and −0.437 (−0.582 to −0.406; q = 9.93e-09) (S2 Table). These are changes in mean log2 relative expression across set members, not UCell units. The paired edgeR gene-level analysis further showed that a signature-level estimate should not be interpreted as uniform regulation of every SG component (S4 Fig). S6 Data provides the complete gene-level tables, with missing inferential values for genes failing expression filtering, together with the SG27 subset, filter records, model sample sizes, normalization factors and raw pseudobulk count matrices.

In the post-result comparison using identical depth-retained cells and weights, GSE178341 gene-Z SG-A and SG-B contrasts were 0.040 (0.005 to 0.093; q = 0.0658) and 0.043 (0.014 to 0.072; q = 0.112), respectively, whereas the corresponding UCell contrasts were negative. The gene-Z diagnostic estimates did not meet the eight-test BH threshold. This direction difference was not explained solely by donor eligibility, but did not demonstrate that either score was biologically correct.

### Sorted fractions and spatial regions differed in compartment contrasts

In the six-patient FACS dataset, SG-A FAP-minus-EPCAM expression averaged 0.081 log2 units (95% CI −0.133 to 0.315; q = 0.688); none of the four sets reached the BH threshold. This estimate was imprecise and did not establish equivalence. In GeoMx WTA data, stromal-minus-epithelial SG-A expression was lower in both core and invasive-front regions: −0.357 (−0.456 to −0.260) and −0.339 (−0.396 to −0.281), respectively, among eight paired T1 patients (both q = 0.00781). Among nine patients, invasive-front-minus-core SG-A was −0.106 (−0.170 to −0.047; q = 0.0268) in PanCK-positive segments and −0.113 (−0.204 to −0.006; q = 0.0742) in PanCK-negative segments. These were mixed stromal regions, not purified CAFs. Ribosomal and housekeeping reference changes also differed by compartment, limiting SG-specific interpretation (S7–S8 Data).

### CosMx contrasts depended on fibroblast subtype

The verified CosMx object contained 1257627 QC-passed cells, 6175 measured features and 11 patients. Coverage was only 5/12 for SG-A, 11/27 for SG-B, 2/4 for core and 3/8 for nucleators; G3BP1, TIA1, CAPRIN1 and UBAP2L were absent. All four lists therefore failed the predefined coverage gate. We did not impute absent genes or create a replacement activity score. The 11 measured members were G3BP2, TIAL1, ATXN2, DDX3X, USP10, DDX6, EIF4E, EIF4G1, EIF4A1, EIF4A2 and STAU1.

Author labels included 20201 trophocyte-like CAFs, 2726 trophocytes and 12304 telocytes. Ten patients met the paired 20-cell threshold. No gene reached q <0.05 for trophocyte-like CAFs versus trophocytes before or after depth balancing, or among the eight patients retained by the label-confidence sensitivity. G3BP2’s raw mean patient difference was −0.0064 (95% CI −0.0544 to 0.0384; q = 0.793), changing to 0.0007 (−0.0411 to 0.0397; q = 0.977) after depth balancing. Against telocytes, ten genes had negative raw contrasts with BH support and five retained support after depth balancing: DDX3X, EIF4A2, EIF4G1, STAU1 and TIAL1. Thus, the comparison depended on the reference fibroblast subtype (Fig 5a,b). The author k8 assignments covered 39394 cells; we report patient-level cluster composition and expression without assigning SG-defined identities. The fibroblast subtype labels (trophocyte, trophocyte-like CAF, telocyte, MMP CAF) follow the nomenclature assigned by the source study [23] and were used as provided.

**Fig 5.**
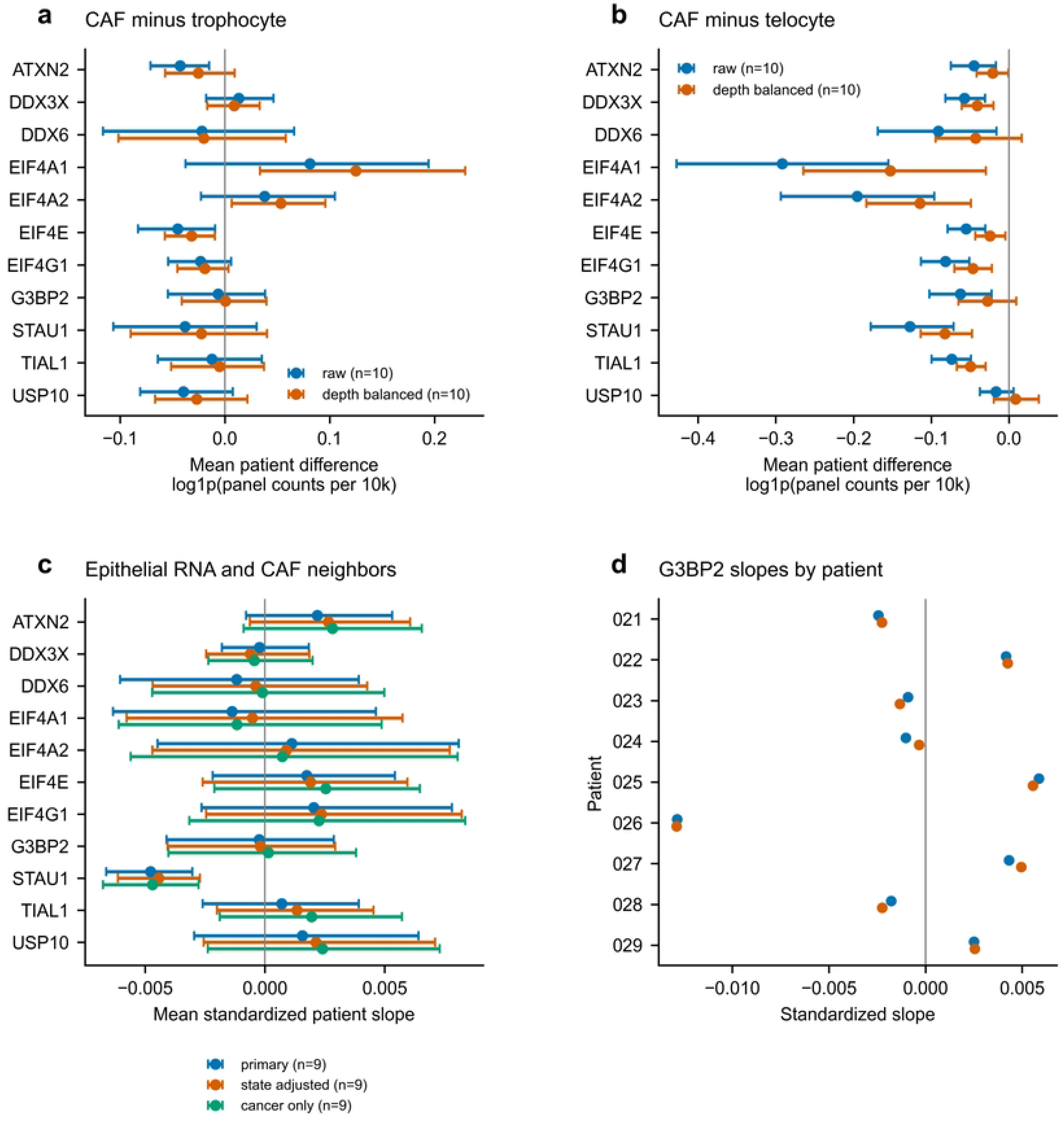
Measured SG-associated genes in CosMx fibroblast states and epithelial neighborhoods. a,b, Equal-weight mean patient contrasts for trophocyte-like CAFs versus trophocytes or telocytes, before and after within-patient depth balancing (ten patients). Bars are 95% percentile intervals from 10,000 patient bootstraps. c, Mean standardized epithelial-gene slopes against 50-µm CAF neighbor fraction, adjusted for panel counts and FOV, with epithelial-state and cancer-related-epithelium sensitivities (nine patients). d, G3BP2 patient slopes for the primary (blue) and state-adjusted (orange) models; G3BP2 is displayed as the measured G3BP-family member, not selected by statistical significance. The 11 measured genes constitute 11/27 SG-B members; no complete SG signature is shown. BH correction covers 11 genes separately per specification. CAF in panels a,b denotes the author trophocyte-like CAF label; in c,d the exposure combines trophocyte-like CAF and MMP CAF fractions. No granule measurements were available.

Of 523346 epithelial cells, 462136 matched the author neighborhood table and 462015 had nonzero neighbor composition. CAF-fraction variation permitted the main model in nine patients comprising 409938 epithelial cells; two patients lacked usable variation. G3BP2’s mean standardized patient slope was −0.00024 (95% CI −0.00411 to 0.00287; q = 0.918). STAU1 showed a small negative slope of −0.00477 (−0.00662 to −0.00303; q = 0.0430), retained after epithelial-state and non-SG-depth adjustment. Leave-one-patient-out means remained negative (−0.00528 to −0.00420). However, the cancer-related epithelial restriction gave −0.00470 (−0.00676 to −0.00277; q = 0.0859), and no gene reached the BH threshold when total fibroblast fraction replaced CAF fraction (11 patients). The neighborhood associations therefore depended on the epithelial population and fibroblast exposure definition (Fig 5c,d; S7 Data).

### Protein correspondence and growth phenotypes varied among SG-list genes

Across 96 matched CPTAC tumors, 11 of the 27 SG genes had positive RNA–protein correlations with BH support; ten retained support after purity adjustment. The median gene-wise correlation was 0.192, with substantial heterogeneity: G3BP1 rho = 0.318 (95% CI 0.122–0.491; q = 0.00436), CAPRIN1 rho = 0.474, STAU1 rho = 0.775 and EIF4G1 rho = −0.006 (Fig 6a). RNA–protein correspondence varied substantially among genes in these bulk tumors. Published U2OS temporal profiles covered 16/27 SG-B members, split into four early, four late and eight profiles labeled Ivariable by the source authors. This descriptive mapping concerned bait-associated protein profiles in heat-stressed U2OS cells, not newly measured CRC dynamics.

**Fig 6.**
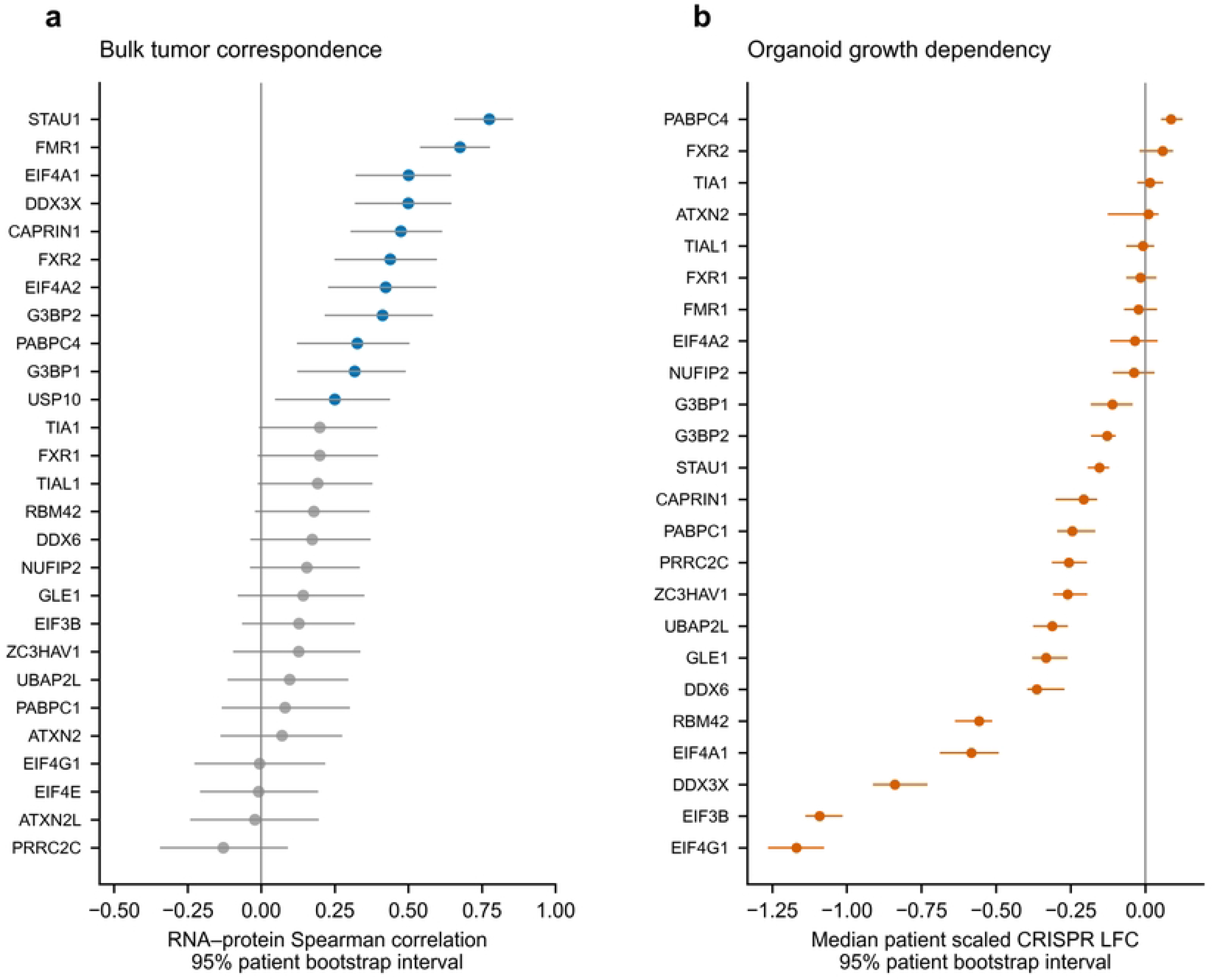
Gene-specific protein correspondence and organoid growth dependency. a, RNA–protein Spearman correlations in prospective CPTAC tumors; blue denotes BH q <0.05 across all 27 genes, gray otherwise. Most genes have 96 matched pairs, with GLE1 at 80 and FXR2 at 92. b, Median across 81 patients of patient-averaged scaled CRISPR log-fold changes from 83 malignant CRC models. All 24 assayed SG-B genes are shown; ATXN2L, USP10 and EIF4E were unassayed. Bars show 10,000-patient-bootstrap 95% intervals; these are pointwise, not simultaneous intervals, and no organoid significance marker is used. Lower scaled log-fold changes indicate stronger growth effects, not more SG-specific dependency. These independent endpoint types are not plotted on a common effect-size scale.

The organoid analysis retained 83 malignant CRC models from 81 patients. Author depletion calls were present in 8/83 models for G3BP1, 1/83 for G3BP2, 0/83 for TIA1 and 1/83 for TIAL1, compared with 58/83 for DDX3X, 73/83 for EIF4G1 and 67/83 for EIF3B. The median patient-scaled log-fold change was −0.110 (95% CI −0.183 to −0.043) for G3BP1, −0.840 (−0.912 to −0.729) for DDX3X, −1.169 (−1.264 to −1.076) for EIF4G1 and −1.091 (−1.139 to −1.014) for EIF3B (Fig 6b). Such growth dependencies can reflect general translation or other functions; they do not demonstrate SG-mediated vulnerability. Forty-seven malignant CRC models overlapped HCMI, and those resources were not treated as independent validation. Missing clinical survival times in CPTAC and only eight outcome events in the locally available HCMI expression-tissue subset precluded a prognostic model (S8 Data).

## Discussion

Paired analyses identified lower fibroblast relative expression of SG-A and SG-B under the pseudobulk definition in both CRC cohorts, and lower SG-B scores under the UCell definition. The direction was less stable across scoring and depth conditions. In the second cohort, gene-Z and UCell estimates differed in sign even when cells, donors, bins and weights were identical; the positive gene-Z diagnostic estimates did not meet their eight-test BH threshold. The study thus identifies reproducible expression contrasts and specifies the scoring and depth conditions under which their direction changes.

Organoid growth data provide a functional context for this distinction. Depletion calls were frequent for DDX3X, EIF4G1 and EIF3B (58/83, 73/83 and 67/83 models), compared with G3BP1, G3BP2, TIA1 and TIAL1 (8/83, 1/83, 0/83 and 1/83). This is a descriptive comparison of selected genes, without a test of functional-category enrichment. The screens measured growth under their assay conditions rather than granule assembly under defined stress, and lack of a depletion call does not establish dispensability for stress-induced granule formation. The high-depletion genes also have translation-associated functions important for growth. The contrast therefore supports interpreting the lists as mixtures of functional roles and helps explain why an aggregate transcript score can reflect processes beyond granule assembly.

The scoring approaches summarize different quantities. Gene-Z gives each gene a scale relative to the analyzed cell population; UCell describes its position within each cell’s expression ranking; pseudobulk log-CPM describes relative abundance in an aggregated RNA pool. Agreement across these quantities strengthens an expression comparison, whereas disagreement identifies a dependence on the measurement definition. The paired gene-level results show heterogeneity among set members as well. Reproducibility is consequently most informative when the quantity summarized is stated explicitly and held constant across cohorts.

Depth sensitivity was a substantial component of the comparison. Library size reflects capture efficiency, RNA content and cellular state, so normalization or regression can change the biological contrast as well as reduce technical imbalance [28,29]. Within-donor balancing selects cells with overlapping depth distributions and is a sensitivity analysis, not a causal estimate of technical sequencing depth. It reduced observed depth differences, while same-donor comparisons showed that donor attrition alone did not explain the second cohort’s sign change. Reporting both the retained donors and retained cells makes the population underlying each estimate explicit.

Detection patterns help explain why set composition matters. The SG lists had higher per-gene detection than SenMayo, yet their smaller sets had appreciable all-zero fractions. An all-zero detection pattern consequently has a different interpretation from absence of the encoded protein or a granule, and droplet sparsity alone is not evidence of a separate zero-inflation process [30]. SenMayo and SASP were not validated controls for the sensitivity or specificity of an SG assay. Their positive contrasts instead provide empirical examples of other compartment-associated expression signals retained by the same analysis.

SG assembly depends on protein interactions, stress conditions and RNA recruitment [1,3,13]. The retrospective catalog comparison reinforces the distinction between encoding an SG-associated protein and having an mRNA classified as SG-enriched in a particular experiment. Neither annotation calibrates an expression score to granule abundance. KRAS-stratified contrasts were not tested here, although experimental links between mutant KRAS and SG regulation [4] suggest a possible source of biological heterogeneity. Joint measurements of genotype, transcript abundance and protein localization could distinguish this source of variation in a future compartment comparison.

The reference measures require similar care. SASP is nested within SenMayo, and secretory or inflammatory expression can reflect fibroblast identity or activation as well as senescence [15,31]. These lists provide empirical comparisons, not independent senescence diagnoses. The housekeeping list also includes VIM and B2M, whose expression can vary by cellular context. Its positive fibroblast contrast is therefore interpretable as a feature of that reference list rather than an invariant technical baseline.

Competitive backgrounds add a separate assessment of specificity [19,20]. An SG list can have a non-zero donor contrast while resembling other genes in the same compartments. This comparison was limited to the sampled candidate pools and observed donors in GSE132465 and did not preserve every aspect of gene–gene correlation. Its matched background was shifted below zero. The result illustrates why a gene-set specificity comparison should use the empirical background distribution rather than assume symmetry around zero.

Spatial and protein measurements refine the questions worth testing next. CosMx showed that fibroblast subtype choice changed gene-level contrasts, while missing panel genes prevented a complete-signature comparison. STAU1 combined a strong RNA–protein correlation with a small CAF-neighborhood association, making it a candidate for targeted follow-up. This nomination is exploratory: the spatial association was sensitive to the epithelial population and exposure definition, and neither bulk protein abundance nor neighborhood composition measures SG recruitment. A useful next test would measure protein localization and dual-marker granule formation alongside transcript abundance in defined fibroblast and epithelial populations.

Previous studies established that signature scoring can be biased in single-cell data [6,7,8], and SG-related CRC outcome models have already been reported [5]. This study adds a donor-paired examination of a fixed compartment comparison, with explicit eligibility, depth sensitivities and complete discordant results. It supports reporting four items together: the scoring definition, donor or sample unit, depth treatment and measured gene coverage. A focused stress-and-recovery experiment with the fixed RNA panel could test temporal correspondence with granule formation; adequately replicated fibroblast and epithelial models would then be needed to address the compartment contrast.

The main limitations concern the analyzed populations and the available measurements. We retained upstream processing and cell annotations and did not repeat raw-read quality control. The cohorts differed in tissue composition, patient characteristics and sequencing context, and broad compartments retained unmodeled cell-state heterogeneity. Some donor means were based on few cells; requiring 20 cells reduced the paired populations to 15 and 29 donors. TMM depends on an appropriate scaling reference, and depth restriction changes the cells represented. CosMx measured 11/27 genes in at most 11 patients and inherited label uncertainty, possible classifier-feature overlap, segmentation error and spatial confounding. External resources used different populations and endpoints and therefore provide complementary context rather than interchangeable replication. The analyses were exploratory, local multiplicity adjustments did not control a project-wide error rate, and no equivalence design or direct granule assay was available.

## Conclusions

Fibroblast–epithelial SG gene-set contrasts in CRC depend on how expression is summarized and how sequencing depth is handled. Paired pseudobulk and selected rank-based contrasts reproduced across cohorts, while comparisons across scoring definitions could differ in direction. Gene coverage, fibroblast reference state and heterogeneous protein and growth phenotypes further specified the interpretation. Reporting the scoring definition, retained donor population, depth sensitivity and measured genes makes these transcript comparisons interpretable and provides a defined starting point for direct granule experiments.

## Data and code availability

The core single-cell studies are deposited under GSE132465 and GSE178341; the sorted fractions are GSE39396. The spatial resource is available at https://doi.org/10.5281/zenodo.17671259; the CosMx experiment-2 archive used here has MD5 8a128cdff99e61873a4b6405925f45f7 and is content-identical in record 20136513. Prospective CPTAC-COAD matrices are distributed at https://www.linkedomics.org/data_download/CPTAC-COAD/. The published SG protein-profile resource is associated with PXD044967 and the supplementary tables of reference [14]; organoid data are supplied with reference [26]. S1–S6 Data preserve the core definitions, donor records, scores, uncertainty estimates, FAP inputs, competitive draws, UCell and pseudobulk results. S7 supplies spatial coverage, patient-level subtype and neighborhood results, exclusions and scripts; S8 supplies sorted-fraction, RNA–protein, protein-profile, organoid and clinical-feasibility records. File inventories, dictionaries and SHA-256 manifests accompany the attachments. The full CosMx RDS and original all-gene scRNA-seq objects are not duplicated in the attachments; cell-level spatial analyses require the identified public source files. Executable patient-summary reconstruction is included. Analysis code and processed reproducibility data are publicly available at https://github.com/wangjinglin0905-gif/sg-crc-transcript-interpretation (version v05.2: https://github.com/wangjinglin0905-gif/sg-crc-transcript-interpretation/releases/tag/v05.2). The permanent archive is available at https://doi.org/10.5281/zenodo.22985583. The retrospective catalog comparison added for this manuscript is provided separately in S3 Table and S9 Data; these additions are not included in the v05.2 archive. The public analysis version remains v05.2.

## Author contributions

Zaili Yang: Writing – original draft, Data curation, Formal analysis. Fei Li: Writing – review & editing, Formal analysis, Investigation. Yin Chen: Methodology, Formal analysis. Yali Shi: Software, Visualization, Validation. Yu Li: Writing – review & editing, Validation, Supervision. Jinglin Wang: Writing – review & editing, Conceptualization, Supervision, Software.

## Supporting information

**S1 Table. Gene-Z comparisons among the same retained donors.** Paired changes distinguish donor selection from within-donor depth sensitivity.

**S2 Table. UCell and pseudobulk contrasts with at least 20 cells per donor and compartment.** Patient-paired estimates, confidence intervals and local multiplicity adjustment.

**S3 Table. Retrospective annotation of the fixed SG gene lists against public resources.** Exact membership and GOA, Khong transcript and Hu protein-profile annotations; source-specific meanings and missing-coverage labels are retained. This is a later catalog comparison, not the original derivation of the lists.

**S1 Fig. Empirical reference contrasts.** Tumor-only unbalanced and depth-balanced estimates for SenMayo, SASP, ribosomal and housekeeping lists, and MKI67. Points and intervals have the same definitions as Fig 1. * The first four measures average gene-wise Z scores; MKI67 uses log-normalized expression. The measures have different biological meanings and are not mutually independent positive controls.

**S2 Fig. Direction and magnitude under five score constructions.** Color indicates only the sign of the tumor-only median donor contrast; numbers report the corresponding estimate. The five constructions use different units, so magnitudes must not be compared as fold changes. Top-five expression is a custom statistic and is not UCell or AUCell.

**S3 Fig. Competitive random-gene-set distributions in GSE132465 tumor tissue.** Panels a–d correspond to SG-A, SG-B, core and nucleator lists. Each outline histogram contains 2,000 random sets of the target size. Matched sets were sampled from stored expression/detection candidate pools; unmatched sets were drawn from the 25,258-gene universe. Dashed lines denote observed median donor contrasts across 23 donors. Tail areas are computed from the empirical distribution without assuming a zero-centered null. The distributions describe competition among gene sets conditional on observed donors, not statistical power across new donor samples.

**S4 Fig. Gene-level components and donor heterogeneity.** a, Paired edgeR quasi-likelihood model log2 fold changes for all 27 SG-B genes; dots indicate BH FDR <0.05 against all filtered genes within each cohort, not just these 27. Color is capped at ±1.2 for readability; exact values are in S6 Data. b, All min20-donor SG-A UCell contrasts ordered separately within each cohort. Ordering is for display and does not match individuals across datasets. Both panels concern relative RNA expression/ranking, not granule measurements.

**S1 Data. Gene definitions, coverage and overlap.** Exact memberships for the four SG and reference measures, cohort-specific missing members and pairwise overlap. Author-curated SG lists are not validated granule-activity signatures.

**S2 Data. Cohorts, tissue strata and donor eligibility.** Cohort and donor-by-tissue cell counts, the GSE178341 patient–tissue mapping, min1/min20 eligibility, pseudobulk sample records and processed-source hashes.

**S3 Data. Gene-Z donor contrasts and uncertainty.** All-tissue, tumor-only, normal-only and tissue-balanced results, donor-level contrasts, 95% bootstrap intervals, local BH adjustment, same-donor and depth-bandwidth sensitivities, shared SenMayo analysis and corrected cell scores.

**S4 Data. Detection and scoring diagnostics.** Mean gene detection, all-zero fractions, depth associations, five score constructions, and matched/unmatched competitive summaries and draws. Random-gene-set intervals are not donor CIs or equivalence limits.

**S5 Data. FAP donor-level comparisons.** Tumor-fibroblast scores and FAP detection labels, patient eligibility and bin support, individual donor contrasts and the 36 FAP summary rows. The supplied reconstruction script preserves the original bootstrap RNG states.

**S6 Data. UCell and paired pseudobulk analyses.** All 64 set-level summaries and donor differences, UCell cell scores, pseudobulk integer counts and set summaries, complete edgeR gene results, SG27 results, filtering, model-sample metadata, official UCell core source and a summary-statistics entry point. Each set-level analysis specification uses an eight-test BH family; gene-level FDR is calculated over all retained genes within each cohort.

**S7 Data. Spatial coverage and patient-level CosMx and GeoMx analyses.** Panel features, coverage failure, frozen specifications and amendments, subtype contrasts, label/depth sensitivities, author k8 descriptions, neighborhood slopes and leave-one-patient-out estimates, spatial region contrasts, source checksums and executable scripts. Full source expression objects are accessed from the public archive.

**S8 Data. External sorted fractions protein context organoid dependency and clinical feasibility.** FACS mapping and patient contrasts; matched CPTAC RNA–protein values and correlations; published SG protein-profile annotations; organoid model selection, patient summaries and overlap audit; and the decision not to build a prognostic model. Protein time profiles are author means, not new replicate-level experiments.

**S9 Data. Gene-catalog annotation sources and reconstruction.** The 27-gene comparison, target-level evidence rows, resource versions, input hashes, dictionary and executable verification script. This manuscript-specific addition is separate from the public v05.2 analysis archive.

